# Nicotinamide Riboside alleviates exercise intolerance in ANT1-deficient mice

**DOI:** 10.1101/2022.02.25.481689

**Authors:** Patrick M. Schaefer, Jessica Huang, Arrienne Butic, Caroline E. Perry, Tal Yardeni, Wendy Tan, Ryan Morrow, Joseph A. Baur, Douglas C. Wallace

**Affiliations:** Center for Mitochondrial and Epigenomic Medicine, Division of Human Genetics, Department of Pediatrics, Children’s Hospital of Philadelphia, PA, 19104; Department of Physiology and Institute for Diabetes, Obesity, and Metabolism, Perelman School of Medicine, University of Pennsylvania, Philadelphia, PA, USA; The Bert Strassburger Metabolic Center, Sheba Medical Center, Israel, 52561; Department of Pediatrics, Perelman School of Medicine, University of Pennsylvania, Philadelphia, PA, 19104

**Keywords:** mitochondrial disorder, exercise, nicotinamide riboside, NAD^+^/NADH

## Abstract

Mitochondrial disorders are often characterized by muscle weakness and fatigue. Null mutations in the heart-muscle adenine nucleotide translocator isoform 1 (ANT1) of both humans and mice cause cardiomyopathy and myopathy associated with exercise intolerance and muscle weakness. We have analyzed the exercise physiology of mice deficient in ANT1, demonstrating a peripheral limitation of skeletal muscle mitochondrial respiration. Upon exercise, lack of Nicotinamide adenine dinucleotide^+^ (NAD^+^) results in a substrate limitation and stalling of the TCA cycle and mitochondrial respiration. Treatment of ANT1-deficient mice with nicotinamide riboside increased NAD^+^ levels in skeletal muscle and improved the exercise capacity and mitochondrial respiration. Thus, increasing NAD^+^ levels with nicotinamide riboside can alleviate the exercise intolerance associated with ANT1-deficiency, indicating the therapeutic potential of NAD^+^-stimulating compounds in specific mitochondrial myopathies.

## Introduction

Adenine nucleotide translocators (ANTs) mediate ADP/ATP exchange across the inner mitochondrial membrane exchanging ATP from the mitochondrial matrix with ADP from the cytosol. In addition, the ANTs play an essential role in mitophagy, provide a voltage sensitive proton channel and regulate the mitochondrial permeability transition pore (mPTP) (1–4).

Humans have three ANT isoforms, which show a tissue-specific expression pattern. Null mutations in the nuclear-encoded heart/muscle isoform ANT1 cause mitochondrial myopathy and cardiomyopathy in humans (5, 6). Certain missense mutations in the human ANT1 gene can manifest as autosomal dominant progressive external ophthalmoplegia (7). ANT1-null patients show elevated serum lactate levels, decreased phosphocreatine levels in muscle and exercise intolerance consistent with an impaired ATP export from the mitochondria. In addition, patients show ragged red muscle fibers and accumulation of aberrant mitochondria due to the defect in mitophagy (5, 6).

Similarly, mice deficient for ANT1 (*Slc25a4^−/−^*) manifest a severe exercise intolerance, cardiomyopathy, ragged red muscle fibers, mitochondrial hyperproliferation, impaired coupled respiration, and increased uncoupling of skeletal muscle mitochondria (8, 9). In addition, glucose homeostasis is altered in ANT1-deficient mice, which manifests as an improved insulin-sensitivity, glucose tolerance and resistance to a high-fat diet (10, 11). In summary, the ANT1-deficient mouse model mirrors the human pathology with respect to myopathy, cardiomyopathy and exercise intolerance (9). Thus, exercise intolerance in ANT1-deficient mice could serve as a marker for muscle dysfunction in humans.

It is known that NAD^+^ levels decrease with age and are lower in many disorders with an underlying mitochondrial dysfunction (12, 13). Boosting NAD^+^ levels with compounds such as nicotinamide riboside (NR) or nicotinamide mononucleotide (NMN) can be beneficial in a range of metabolic disorders (reviewed in (14)), including neurodegeneration (15), obesity (16) and diabetes (17). Similarly, NAD^+^-boosting compounds have been shown to rescue exercise intolerance and mitochondrial myopathy in mouse models of mitochondrial disease and patients with adult-onset mitochondrial myopathy (18–20). In the current study, we delineate the physiological limitations underlying the exercise intolerance in ANT1-deficient mice. We then show that NR can alleviate the associated exercise intolerance.

## Results

### VO_2_ is decreased in ANT1 mice during exercise, contributing to their exercise intolerance

We assessed the exercise capacity in C57Bl/6J mice and *ANT1^−/−^* (*Slc25a4^−/−^*) mice (called “B6 control” and “ANT1” from here on) ranging from 24 weeks until 70 weeks of age. As observed previously, ANT1 mice show a lower VO_2max_ and a shorter time until exhaustion in the ramp test on a metabolic treadmill by 6 months of age (Fig.1a). In contrast to B6 control animals, ANT1 mice reach their VO_2max_ shortly after the start of the exercise stress test and subsequently their VO_2_ declines despite an increasing exercise intensity (Fig.1a, blue line). In B6 control mice, VO_2_ steadily increases with exercise intensity, reaching VO_2max_ upon exhaustion (Fig.1a, black line).

**Figure 1:**
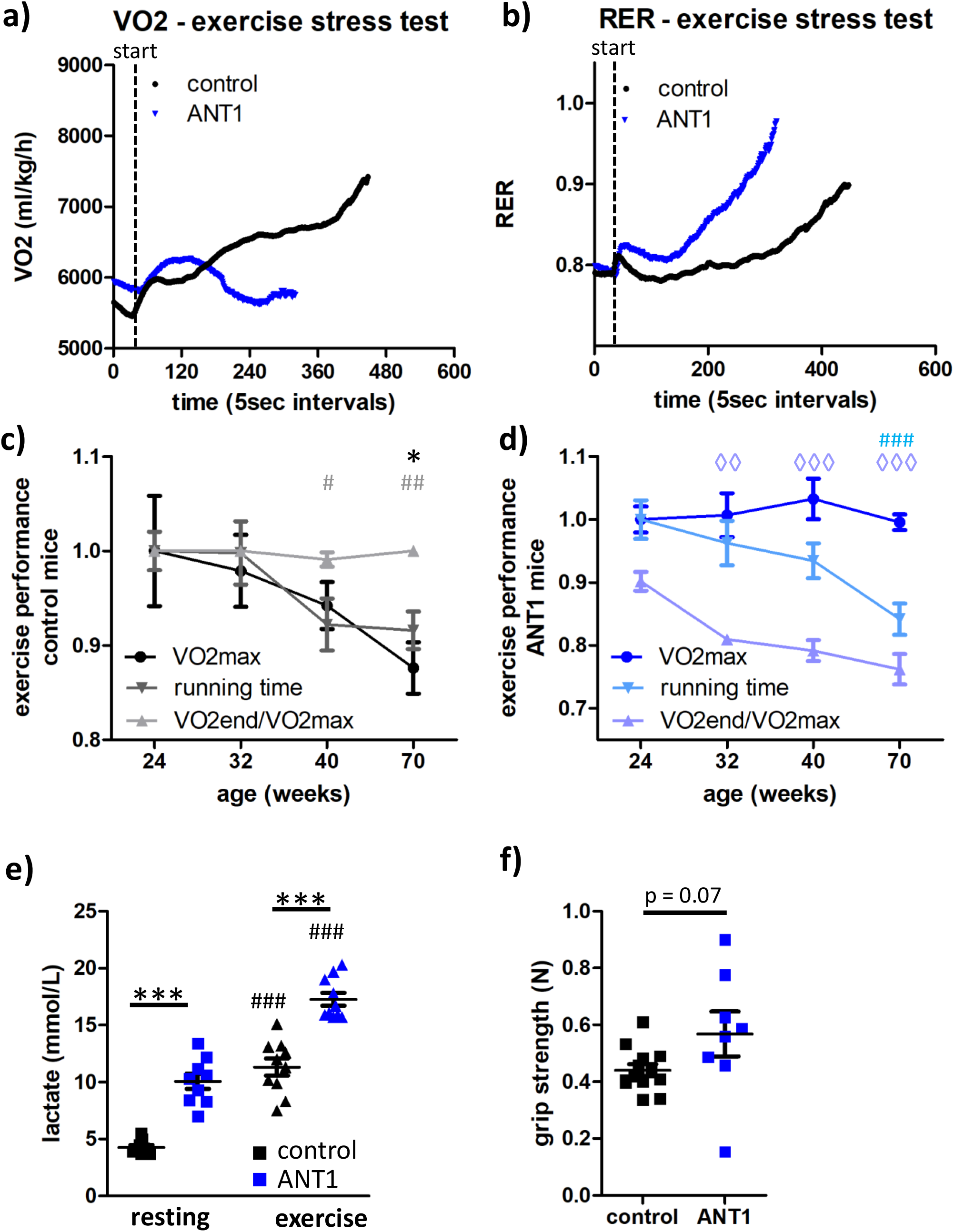
VO_2_ decreases in ANT1 mice during exercise. **a/b)** Kinetics of the average oxygen uptake (VO_2_) (a) and respiratory exchange ratio (RER, VO_2_/VCO_2_) (b) of 4-month-old mice during an exercise stress test (ramp protocol) on a metabolic treadmill (n = 24-40, curve displayed until >50% of the animals quit). **c/d)** Display of exercise performance parameters (VO_2_max, running time, VO_2_end/VO_2_max) in B6 control mice (c) and ANT1 mice (d) over age normalized to performance at 24 weeks (n = 7-9, error bars indicate SEM). Significances were calculated compared to performance at 24 weeks of age and are indicated as * for VO_2_max, # for running time and ◊ for VO_2_endVO_2_max. **e)** Blood lactate levels in resting mice and immediately after exhaustive exercise (n = 9-10, 6-9-month-old). **f)** Average grip strength of front paws of 18-month-old B6 control and ANT1 mice (n = 8-13, average of 3 trials/mouse). Significances between strains (*) and between “exercise” and “resting” (#) were calculated using Mann Whitney test (*p < 0.05, **p < 0.01, ***p < 0.0001).

B6 control mice show a relatively steady respiratory exchange ratio (RER) of 0.8 for the first two-thirds of the exercise stress test and a subsequent increase, indicating an increased carbohydrate usage upon high exercise intensity (Fig.1b). In contrast, the RER increases very fast in ANT1 mice starting at the time point of VO_2max_ (Fig.1b). This demonstrates that reduced oxygen uptake coincides with a fast switch from fat to predominantly carbohydrate fuel usage in ANT1 mice, likely due to increased glycolytic energy production. As expected, VO_2max_ and running time (time until exhaustion in the exercise stress test ramp protocol) decrease with age in B6 control mice (Fig.1c), as does running time in ANT1 mice (Fig.1d). However, VO_2max_ does not decrease in ANT1 mice with age, although the VO_2_ decrease upon running becomes more exaggerated, demonstrated by calculating a ratio of the VO_2_ at exhaustion (VO_2end_) and the VO_2max_. Thus, in ANT1 mice VO_2end_/VO_2max_ correlates with the running performance over age (Fig.1d), suggesting that decreasing VO_2_ during exercise is an important factor in the exercise intolerance of ANT1 mice.

To further understand the decreasing VO_2_ during endurance exercise in ANT1 mice, we first tested for a psychological or motor limitation, which could prevent the ANT1 mice from exhausting themselves physically in the exercise stress test. Maximal RER is used as a measure of exhaustion with RER >1.0 indicating exhaustion (21). Both ANT1 and B6 control mice show an average RER_max_ of >1.0, with ANT1 mice displaying a longer recovery period for RER and VO2 to return to baseline following an exercise stress test (Fig.S1a-c).

We measured blood lactate levels both at rest and immediately after an exercise stress test. ANT1 mice showed higher lactate levels at rest and after exercise compared to B6 control mice, resulting in a similar increase in blood lactate levels upon exercise (Fig. 1e). Thus, ANT1 mice show higher tolerance of exhaustion compared to B6 control mice. Furthermore, we tested motor coordination in 18-month-old ANT1 and B6 control mice and found a similar performance on the rotarod (Fig.S1d), indicating no motor phenotype that could explain a reduced exercise performance on the treadmill. Similarly, grip strength was not reduced in 18-month-old ANT1 mice compared to control (Fig.1f), suggesting no deficit in muscle strength. In summary, we could demonstrate a reduced exercise capacity in ANT1 mice due to physical limitations.

### Skeletal muscle oxygen consumption limits VO_2_ in ANT1 mice

Physical factors limiting VO_2max_ can be divided into a central limitation by the cardiovascular system and the transport of oxygen to the muscles or peripheral limitation by the mitochondrial oxygen consumption of the muscles. We assessed whether ANT1 mice are centrally or peripherally limited in their VO_2_.

First, we performed an exercise stress test at 100% oxygen, thereby increasing oxygen availability. This resulted in an increase in running time in B6 control mice but no change in ANT1 mice compared to running under normoxia. This shows a central limitation in B6 control mice but a peripheral limitation in ANT1 mice (Fig.2a). Next, we compared VO_2max_ reached in the exercise stress test to VO_2max_ reached during cold stress. Both stressors resulted in a similar VO_2max_ in B6 controls but cold stress-induced a higher VO_2max_ in ANT1 mice (Fig.2b). This also indicates a peripheral limitation in ANT1 mice. This is further supported by venous blood gas analysis performed of mice at rest or immediately after an exercise stress test. While there is no difference in the venous partial oxygen pressure (pO_2_) between B6 control and ANT1 mice at rest, ANT1 mice showed a significantly higher venous pO_2_ after exercise (Fig.2c). This indicates that ANT1 mice have normal availability of oxygen to the muscle but insufficient oxygen extraction by the mitochondria. This is in line with no alterations in blood hemoglobin levels in ANT1 mice compared to B6 controls (Fig.2d). Lastly, we measured muscle pO_2_ directly in the gastrocnemius muscle of anesthetized mice. At baseline, muscle pO_2_ was similar between B6 control and ANT1 mice (Fig.2e).

**Figure 2:**
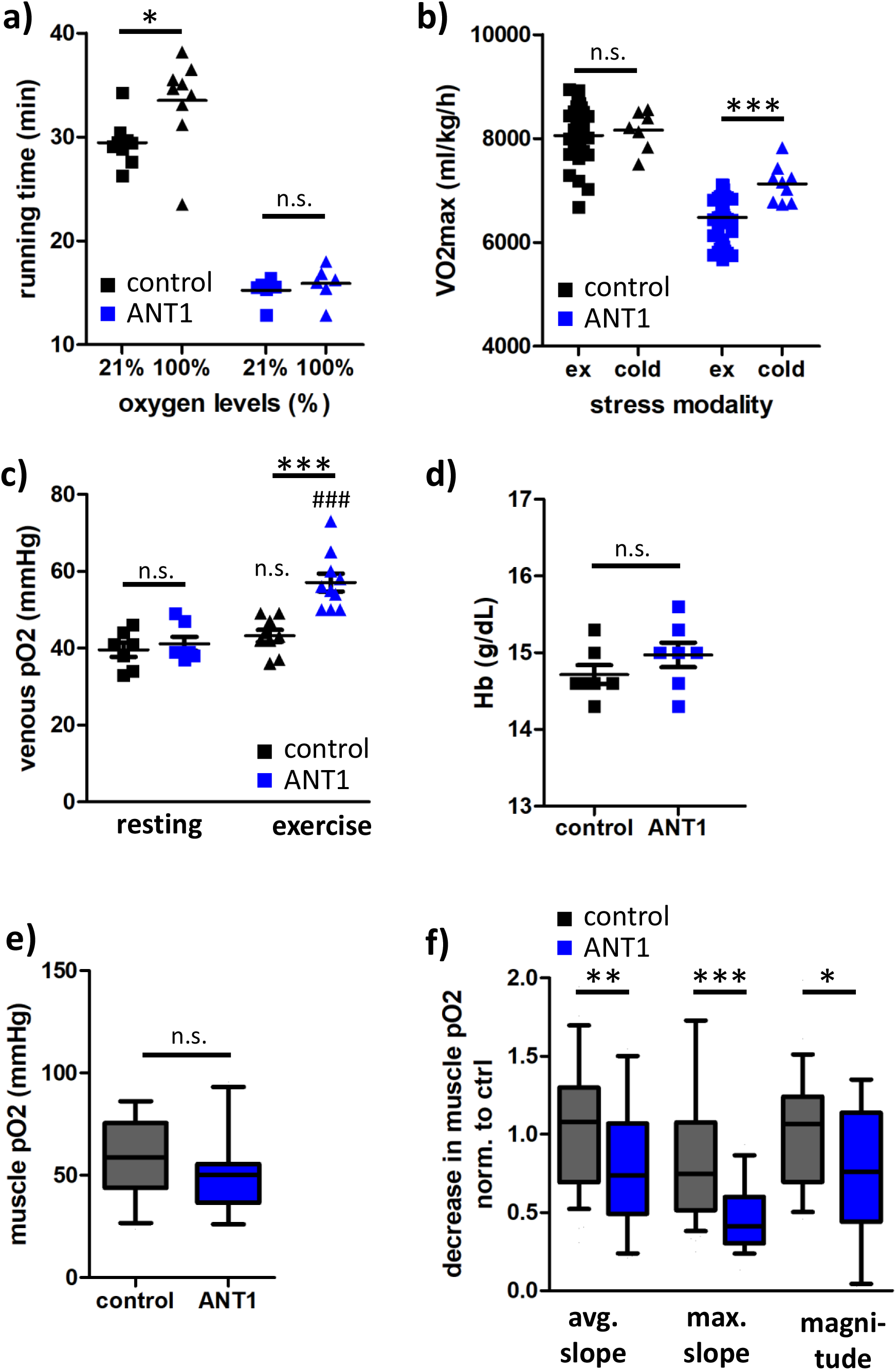
Skeletal muscle oxygen consumption limits VO_2_ in ANT1 mice. **a)** Time until exhaustion in an exercise stress test (ramp protocol) performed at normoxia (21% oxygen) or hyperoxia (100% oxygen) in the same mice (n = 6-9). Significances between normoxia and hyperoxia were calculated using Wilcoxon signed rank test. **b)** VO_2_max reached in an exercise stress test (ex, n = 39-44) or during a 4h cold exposure (cold, n = 7-9, 4°C) in 4-month-old mice. Significances between stress modalities were calculated using unpaired t-test. **c)** pO_2_ in blood from the submandibular vein in resting mice and immediately after exhaustive exercise (n = 7-10, 6-9-month-old). Significances between strains (*) and between “exercise” and “resting” (#) were calculated using Mann Whitney test. **d)** Hemoglobin levels in blood from the submandibular vein in resting mice (n = 7, 6-9-month-old). **e/f)** Muscle pO_2_ in non-stimulated M. gastrocnemius of anesthetized mice (6-12-month-old) (e). Magnitude and rate of decrease in muscle pO_2_ (g) upon sciatic nerve stimulation normalized to B6 controls (n = 17-18, 3 technical replicates). Whisker plots show 10-90% intervals, significances between strains were calculated using Mann-Whitney test.

However, after stimulation of the sciatic nerve, muscle pO_2_ in ANT1 mice decreased significantly less than in control mice (Fig.2f, magnitude) and at a lower average and maximal rate (Fig.2f, avg. and max. slope). This demonstrates that muscle oxygen consumption upon contraction is reduced in ANT1 mice.

In summary, using multiple lines of evidence, we could demonstrate that skeletal muscle oxygen consumption limits the VO_2_ in ANT1 mice.

### NAD^+^ availability stalls TCA cycle and limits skeletal muscle respiration in ANT1 mice upon exercise

To assess oxygen consumption in skeletal muscle directly, we performed high-resolution respirometry of the soleus muscle dissected from non-exercised (resting) mice. Surprisingly, we found no reduction in the oxidative phosphorylation (OxPhos)-capacity (coupled respiration with excess complex I (CI) and complex II (CII) substrates, Fig.3a) and even an increase in the ETS capacity (uncoupled respiration, Fig.3b) in the ANT1 mice compared to B6 control when respiration rate was normalized to tissue mass. Thus, mitochondrial respiratory capacities appear not to limit endogenous respiration.

**Figure 3:**
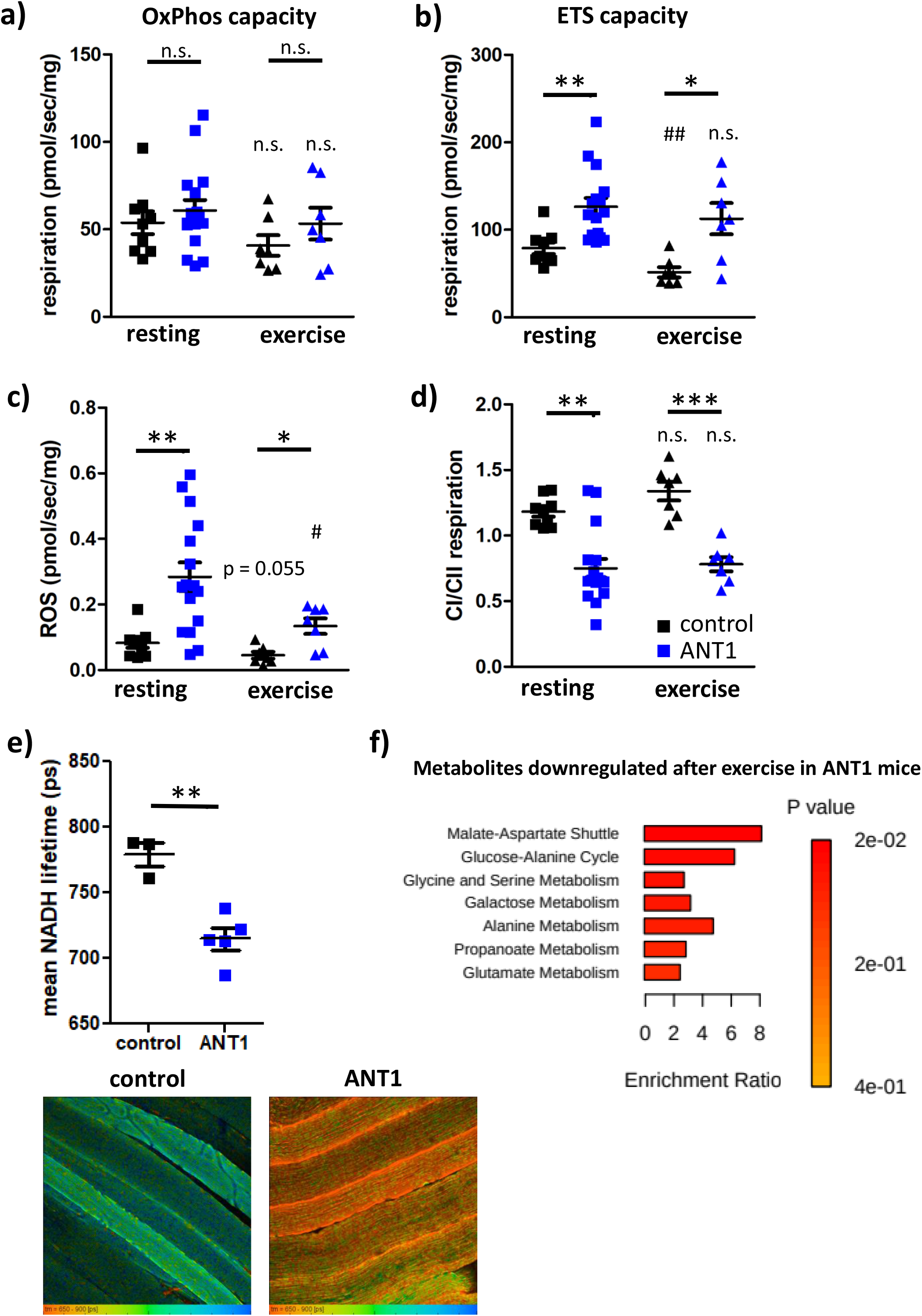
Reduced Complex I (CI)-linked respiration alters redox state and causes substrate limitation in ANT1 mice. **a/b/c)** Coupled (OxPhos capacity, (a)) and uncoupled (ETS capacity, (b)) respiration and ROS production (c) in the presence of complex I and II substrates normalized to tissue mass in M. soleus of 6-month-old mice (n = 7-16) that were at rest or acutely exercised until exhaustion before being sacrificed. **d)** Ratio of CI to CII respiratory capacities in soleus of 6-month-old mice (n = 7-16) that were at rest or acutely exercised until exhaustion before being sacrificed. Significances between strains (*) and between resting and exercise (#) were calculated using Mann Whitney or unpaired t-test (*p < 0.05, **p < 0.01, ***p < 0.0001). **e)** Mean NADH lifetime measured by fluorescence lifetime imaging microscopy of NADH autofluorescence in M. gastrocnemius of 6-month-old mice (n = 3-5, 10 images each). NADH lifetime is false-color coded with red indicating a shorter NADH lifetime (more reduced NAD^+^/NADH redox state) and blue indicating a longer NADH lifetime. f) Enrichment analysis of global metabolomics (downregulated metabolites, p < 0.05) in M. gastrocnemius of resting ANT1 mice compared to ANT1 mice exercised until exhaustion using Metascape. The enrichment ratio is indicated by the length of the bar and the p-value by color.

We found a strongly increased ROS production in skeletal muscle of ANT1 mice compared to B6 control (Fig.3c). Thus, we hypothesized that excessive ROS production upon exercise might impair iron-sulfur clusters of the electron transport system, resulting in lower skeletal muscle respiration and the decreasing VO_2_ upon exercise. To check this hypothesis, we ran mice until exhaustion, sacrificed them immediately afterwards and performed respirometry on soleus muscle. However, we did not observe a significantly lowered muscle respiration in acutely exercised ANT1 mice compared to muscle from resting mice (Fig.3a,b). Surprisingly, we found a significantly lower ROS production in skeletal muscle of acutely exercised ANT1 mice, indicating ROS-mediated inactivation of ETS enzymes did not occur upon exercise.

Interestingly, ANT1 mice showed a significantly lowered ratio of CI-linked to CII-linked respirational capacities (Fig.3d). We performed NADH fluorescence lifetime imaging microscopy (FLIM) of skeletal muscle and revealed a significantly shorter NADH lifetime, corresponding to a more reduced NAD^+^/NADH redox state and a lower CI-linked respiration in ANT1 mice compared to B6 controls (Fig.3e). Given our ANT1 mice but not B6 controls also harbor the complex I variant ND5 m.12352C>T (ND5^S204F^), we created additional control mice, called ND5 mice from here on out, harboring the ND5 m.12352C>T (ND5^S204F^) variant. However, ND5 mice do not display a reduced VO_2max_ or running time, VO_2_ does not decrease during exercise (Fig.S2a) and the CI to CII respirational capacities are similar to B6 controls. Thus, we conclude that the ANT1 mutation and not the ND5 variant is responsible for the phenotype of the ANT1 mice.

If mitochondrial capacities do not limit endogenous respiration, we hypothesized a substrate limitation. To further explore this hypothesis, we performed global metabolomics of gastrocnemius muscle of resting and acutely exercised mice. In B6 control mice, pathway analysis revealed an upregulation of TCA cycle, malate-aspartate shuttle, urea cycle and ammonium recycling immediately post-exercise (Fig.S3a). In ANT1 mice, TCA cycle metabolites, urea cycle and ammonia recycling were also upregulated post-exercise (Fig.S3b); however, the malate-aspartate shuttle and the glucose alanine cycle were downregulated (Fig.3f). A closer look at TCA cycle metabolites revealed a significant upregulation of most metabolites upon exercise both in B6 controls and ANT1 mice. However, α-ketoglutarate is significantly downregulated upon exercise in ANT1 skeletal muscle. This could indicate a lower activity of the isocitrate dehydrogenase, the first step in the TCA cycle requiring NAD^+^, which might be lacking due to the lower CI-linked respiration (see Fig.3d). This is in line with a downregulation of the malate-aspartate shuttle in ANT1 skeletal muscle (Fig.3f). To maintain cytosolic NAD^+^ levels and the essential glycolytic flux in ANT1 skeletal muscle, lactate is formed (see Fig.1e) and the Cori cycle might be preferred over the glucose-alanine cycle (Fig.3f), which would preserve the reducing equivalents and thus not recover NAD^+^. Quantification of nucleotide levels in the gastrocnemius muscle revealed a trend towards a reduced NAD^+^/NADH ratio in ANT1 mice after exercise compared to resting ANT1 mice (Fig.S3d). Surprisingly, absolute NAD^+^ levels were increased significantly in ANT1 mice after exercise compared to resting mice (Fig.S3e), suggesting an increase in the NAD/NADH pool size upon exercise in ANT1 mice. This is in line with a reduction in NAM levels upon exercise in ANT1 mice (Fig.S3f) indicating an increase in the NAD^+^ salvage pathway upon exercise, possibly to counteract a lack of NAD^+^. In contrast, we found no changes to the nucleotide levels in B6 control mouse skeletal muscle upon exercise (Fig.S3d-f).

In addition, global metabolomics revealed an upregulation of methylhistidine metabolism in ANT1 mice upon exercise (Fig.S3b), indicating an increased protein catabolism. Combined with a downregulated ammonium ion export via the glucose-alanine cycle (Fig.S3b) this might result in ammonium accumulation in skeletal muscle of ANT1 mice upon exercise, which was shown to further impair mitochondrial respiration (22).

In summary, we showed that complex I respiration is reduced in skeletal muscle of ANT1 mice. This results in a lack of NAD^+^ to maintain the essential glycolytic flux, TCA cycle flux, and ammonia recycling during exercise.

### Nicotinamide Riboside (NR) alleviates exercise intolerance in ANT1 mice

If NAD^+^ availability in skeletal muscle is the limiting factor in ANT1 mice during exercise, boosting NAD^+^ levels should improve exercise capacity in ANT1 mice.

To test this hypothesis, we treated ANT1 and B6 control mice with 300 mg/kg _body weight_ of NR orally with the food, starting at 4 months of age. NR treatment significantly improved the running time of ANT1 mice but not B6 controls (Fig.4a). In addition, NR increased the VO_2end_/VO_2max_ (Fig.4b) in ANT1 mice, indicating a smaller drop in VO_2_ during running (Fig.4d). While VO_2_ at rest (Fig.S4a) or VO_2max_ (Fig.S4b) were not affected significantly. NR treatment increased the VO_2_ reserve capacity (Fig.S4c) in both ANT1 and B6 control mice. Consistent with an increased glycolytic flux, NR treatment also increased both the RER at rest (Fig.S4d) and during exercise (Fig.4c) in ANT1 mice. In the comprehensive lab animal monitoring system (CLAMS), B6 control but not ANT1 mice showed a higher activity level (Fig.S4e) and running wheel activity (Fig.S4f).

**Figure 4:**
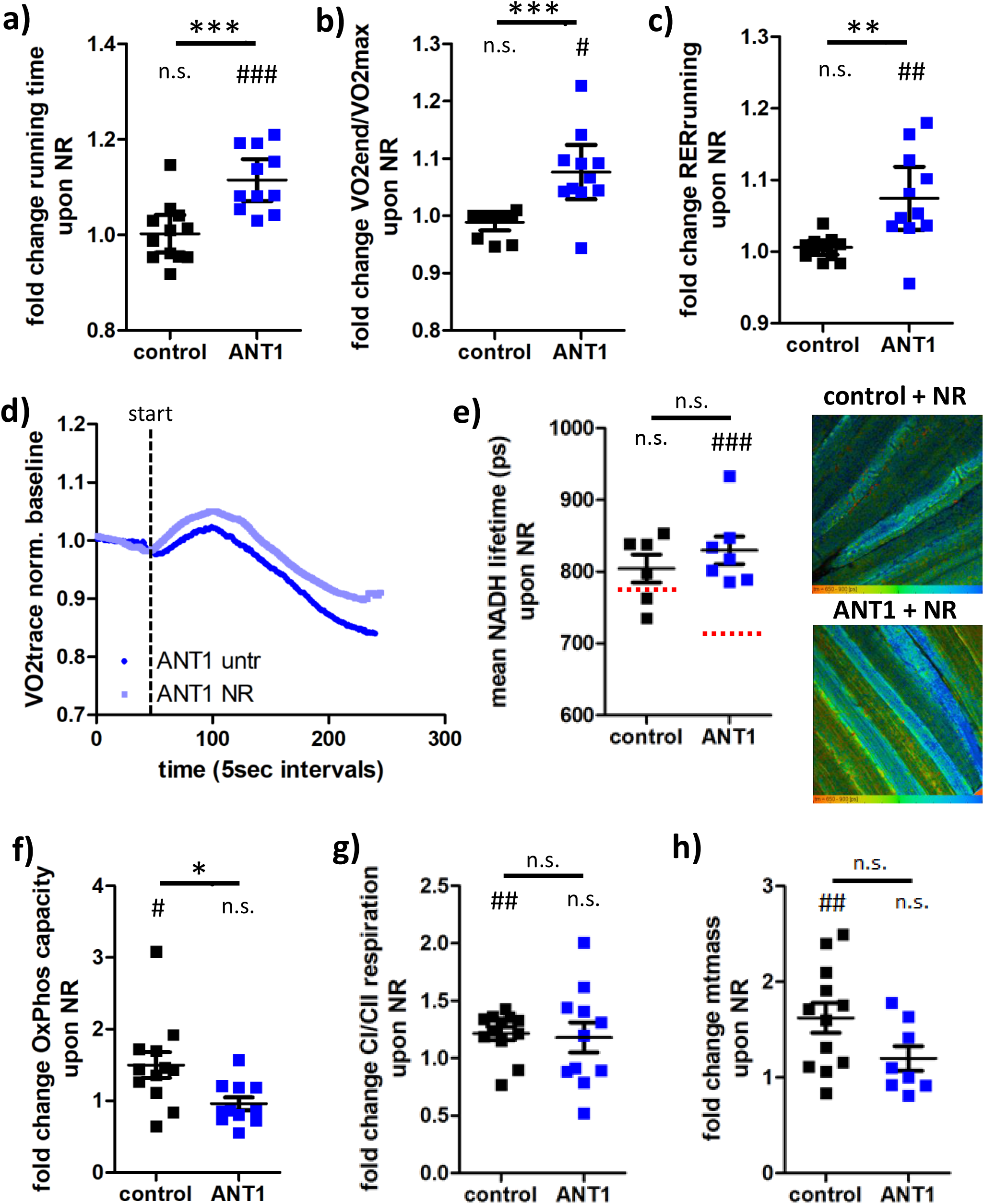
Nicotinamide riboside (NR) alleviates exercise intolerance in ANT1 mice. **a/b/c)** Fold change in running time (a), VO_2_end/VO_2_max (b) and RER (c) in an exercise stress test upon an 8-week treatment with NR in 6-month-old B6 control and ANT1 mice (n = 11-12). d) Average VO2 trace of an exercise stress test in 6-month-old ANT1 mice untreated (untr, dark blue) or treated with NR for 8 weeks (NR, light blue). **e)** Mean NADH lifetime in M. gastrocnemius of B6 control and ANT1 mice treated with NR (n = 6-7, 10 images each). NADH lifetime is false-color coded (same scale as for Fig.3e) with red indicating a shorter NADH lifetime and blue indicating a longer NADH lifetime. Dotted red line indicates the average lifetime of M. gastrocnemius of untreated control and ANT1 mice (see Fig.3e). **f/g/h)** Fold change in M. soleus OxPhos capacity (f), CI/CII respirational capacity (g) and mitochondrial mass (h) upon an 8-week treatment with NR in 6-month-old B6 control and ANT1 mice (n = 8-12). Significances between strains (*) and between NR-treated and untreated (#) were calculated using Mann Whitney or unpaired t-test (*p < 0.05, **p < 0.01, ***p < 0.0001).

NR significantly increased the reduced levels of NAD^+^ content in the gastrocnemius muscle (Fig.S3e) without changing the NAD/NADH ratio (Fig.S4g,h). In contrast, neither NAD^+^ levels nor NAD/NADH ratio were altered in the gastrocnemius of B6 control mice. Hence, oral NR treatment is effective for rescuing NAD^+^ deficiency. We also performed NADH FLIM and found NR treatment increased NADH lifetime in the gastrocnemius muscle of ANT1 mice (Fig.4e).

To assess if the NR-effect is directly due to increased NAD^+^ content or due to stimulated mitochondrial biogenesis and respiration, we performed respirometry on skeletal muscle. In B6 controls, we found NR increased OxPhos capacity (Fig.4f), CI/CII-linked respiration (Fig.4g) and mitochondrial mass (Fig.4h), consistent with stimulated mitochondrial biogenesis. However, the increased muscle respiration did not mediate an increased exercise capacity in B6 control mice consistent with their exercise capacity being limited by cardiovascular delivery of oxygen (see Fig.2). In contrast, in ANT1 mice, NR-treatment did not alter mitochondrial respiration (Fig.4f, g) or mitochondrial mass (Fig.4h) suggesting that the beneficial effect of NR on exercise ANT1 null capacity is due to an increased NAD^+^ content.

### Nicotinamide Riboside reduces exercise capacity in ND6 and CO1 mutant mice

Next, we checked if NR treatment would be beneficial in other mouse models for mitochondrial disorders. Mice harboring a mutation in the complex I subunit ND6 (mtDNA *ND6* m.13997G>A, P25L) (23) (ND6 mice) and mice harboring a complex IV mutation (mtDNA *COI m*.6589T>C, V421A) (24) (COI mice), both show a mild exercise intolerance. An 8-week treatment with NR decreased resting VO_2_ (Fig.5a) but did not alter VO_2_max (Fig.5b) in either ND6 or COI mice, indicating that VO_2_ reserve capacity is not a limiting factor in these mice. However, the exercise capacity measured as running time on the ramp protocol <u>decreased</u> significantly with NR in both strains (Fig.5c). Substrate utilization was not affected by NR treatment in ND6 and COI mice, indicated by no change in RER during rest (Fig.5d) or during running (Fig.5e). Interestingly, both ND6 and COI mice show higher activity levels upon NR treatment (Fig.5f), which reached significance in COI mice.

**Figure 5:**
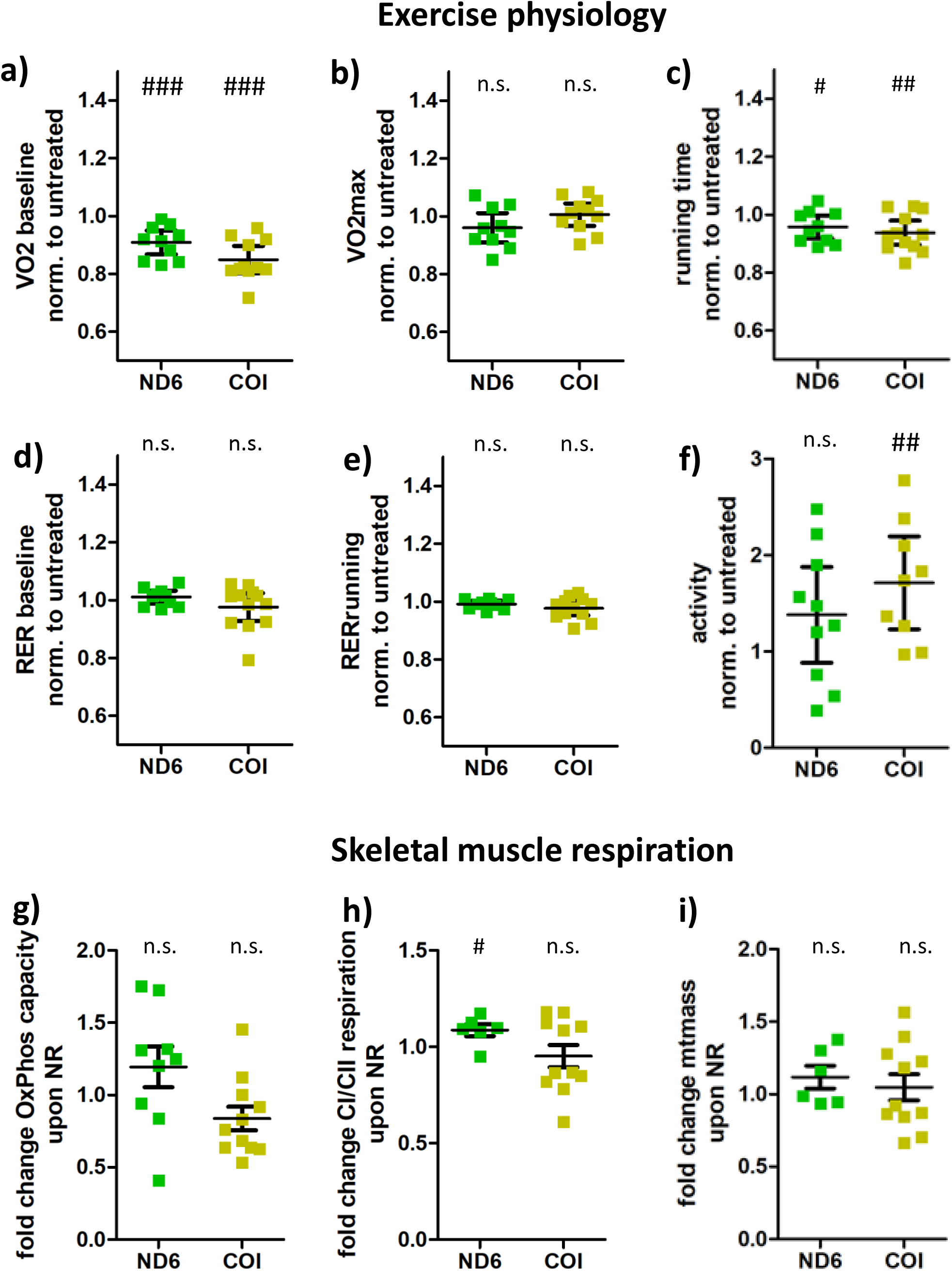
Nicotinamide riboside does not improve exercise intolerance in ND6 or COI mice. **a-e)** Fold change in VO_2_ during rest (a), VO_2_max (b), running time (c), RER during rest (d) and RER during running (e) in an exercise stress test upon an 8-week treatment with NR in 6-month-old ND6 and COI mice (n = 10-12). **f)** Fold change in activity levels during 24h spent in CLAMS upon an 8-week treatment with NR in 6-month-old ND6 and COI mice (n = 9-10). **g-i)** Fold change in M. soleus OxPhos capacity (g), CI/CII respirational capacity (h) and mitochondrial mass (i) upon an 8-week treatment with NR in 6-month-old ND6 and COI mice (n = 6-11). Significances between NR-treated and untreated (#) were calculated using Mann Whitney or unpaired t-test (^#^p < 0.05, ^##^p < 0.01, ^###^p < 0.0001).

Hypothesizing a peripheral limitation to the exercise capacity in the ND6 and COI mice, we checked skeletal muscle respiration in the soleus muscle of NR-treated ND6 and COI mice. Upon NR treatment, ND6 mice show a tendency towards an increased OxPhos capacity (Fig.5g) and a significant increase in CI over CII respirational capacity (Fig.5h). In contrast, COI mice show a tendency towards a lower OxPhos capacity (Fig.5g) and no alteration in CI versus CII respiration (Fig.5h). Similarly, NR treatment did not alter the mitochondrial mass in the soleus muscle of ND6 and COI mice (Fig.5i). These findings are consistent with the absence of an NR effect on VO_2max_ in both strains but do not explain the decrease in running time upon NR treatment. Further experiments on the exercise physiology of ND6 and COI mice would be needed to elucidate the negative effect of NR on the exercise capacity.

We can conclude that NR treatment has a positive effect on mice harboring the nuclear ANT1 mutation due to this mitochondrial defect resulting in reduced NAD^+^ and NAD^+^/NADH ratio. We found that NR treatment did not have a similar beneficial effect for mice harboring mtDNA mutations in the ND6 and COI genes. Hence, the beneficial effects of NR treatment do not seem to be generalizable for all mitochondrial disorders.

## Discussion

Assessing the limiting factors of exercise capacity in ANT1-deficient mice, we gained three crucial insights into how ANT1-deficiency alters exercise physiology and muscle metabolism. First, we discovered unusual VO_2_ kinetics during exercise, with VO_2_ decreasing with increasing workload. We traced this effect to a substrate limitation in the skeletal muscle of NAD^+^. Second, we found that ANT1-deficiency strongly shifts the mitochondrial respiration from CI-linked to CII-linked respiration in skeletal muscle, aggravating the NAD^+^/NADH redox imbalance. Third, we observed that NR can benefit the exercise intolerance in ANT1-deficient mice but did not benefit mice harboring mtDNA ND6 and COI defects.

The VO_2_ kinetics we observed in B6 control mice mirror the VO_2_ kinetics described for humans (25). At moderate exercise levels, the VO_2_ is proportional to the workload (26), thus increasing steadily with increasing running speed (see Fig.1a). At heavy exercise, VO_2_ increases disproportionately to workload due to the addition of the VO_2_ slow component (27), resulting in the steeper rise in VO_2_ towards the end of the exercise regime in B6 control mice. The VO_2_ kinetics at the onset of exercise cannot be assessed in our system because mice are in a metabolic chamber. This causes a delay in detecting changes in the CO_2_/O_2_ ratio relative to activity which is not a concern in the breath-by-breath analysis common in human exercise studies.

In ANT1 mice we could demonstrate that VO_2_ rises initially but then drops with increasing work load, a phenotype that becomes even more prominent with age. We found the VO_2_ is limited by skeletal muscle oxygen consumption in ANT1 mice. This is consistent with pO_2_ decreasing more slowly upon contraction in ANT1 skeletal muscle, indicating a lower endogenous mitochondrial respiration rate. In the ANT1 mice, we found similar respiratory capacities normalized to muscle weight as the B6 control mice, and no indication of a reduced muscle mass, suggesting that the reduced exercise capacity was substrate limited. Based on global metabolomics after acute exercise, respirometry, and rescue with NR we conclude that lack of NAD^+^ might cause a stalling of the TCA cycle and reduction of endogenous respiration. In line with this hypothesis, mice deficient for NAD^+^ due to deletion of Nampt in skeletal muscle show a lower exercise capacity and mitochondrial respiration (28). It would be interesting to see if the Nampt-deficient mice show similar VO_2_ kinetics during exercise as ANT1 mice.

While NR reduced the effect of decreasing VO_2_ during exercise in ANT1 mice, it did not abolish it. Hence NAD^+^ limitation is not the only factor limiting ANT1 muscle performance. ANT1 mice also have an altered glucose homeostasis and we could show that they heavily rely on carbohydrates during exercise, resulting in hypoglycemia, which can cause fatigue (29). Still, pyruvate levels were increased in skeletal muscle of ANT1 mice after exercise, contra-indicating that a lack of glucose impairs respiration. Although we found no reduced respiration and even decreased ROS levels after an acute bout of exercise in ANT1 mice, we cannot exclude a temporary inhibition of iron sulfur clusters by ROS (30) or ammonium ions (22). Tools to assess metabolic flux with a higher temporal resolution would be needed to fully elucidate the skeletal muscle metabolism of ANT1 mice during exercise.

Interestingly, we could demonstrate a shift from CI to CII respiration in skeletal muscle of ANT1 mice, resulting in a more reduced redox ratio. A lack of ATP/ADP exchange would be expected to result in a lower coupled respiration and a higher proton-motive force. One hypothesis for the switch from CI to CII respiration could be that CII does not pump protons, which might favor electrons entering the ETS via CII instead of CI. However, this would not explain why CI capacity is proportionately reduced compared to CII capacity in skeletal muscle of ANT1 mice. A previous study showed an upregulation of all mitochondrial gene expression in skeletal muscle of ANT1 mice, but the data reveal a tendency for a stronger upregulation of CII transcripts compared to CI transcripts (10). Lastly, the higher ROS levels in ANT1 could partially inactivate iron-sulfur clusters of CI. The lower CI to CII respiration results in a more reduced NAD^+^/NADH redox ratio, which contributes to the lack of NAD^+^. Moreover, the NAD^+^/NADH ratio is known to mediate mitochondria-nuclear signaling by affecting histone modifications, which could further contribute to the pathological phenotype of ANT1 mice.

We observed that NR improved the exercise capacity in ANT1 mice but not B6, ND6, or COI mice. These results are consistent with the differential effects of NR on rodent exercise capacity with some studies reporting beneficial effects (31, 32), no effects (19, 33), or negative effects (34). It has been suggested that boosting NAD^+^ is more effective in aged individuals than younger ones (32, 35) due to the decreasing levels of NAD^+^ and NAD^+^ salvage pathway expression with age (reviewed in (13)). Given that NR did not increase NAD^+^ levels in B6 mice, this could explain why we observed no positive effects in this strain. In contrast, ANT1 mice show slightly reduced NAD^+^ levels and a reduced expression of Nampt, possibly due to increased inflammation and oxidative stress (36).

ANT1 null mutations cause myopathy and cardiomyopathy in humans (37). It has been shown that redox imbalance and Nampt deficiency in the heart can contribute to heart failure (38, 39), suggesting a possible role of NAD^+^ deficiency in mediating cardiac remodeling in ANT1 null patients. In contrast, ANT1 missense mutations mostly affect the extraocular muscles (EOM) in humans. Interestingly, lactate dehydrogenase expression levels are low in human EOM compared to other skeletal muscles (40). This indicates that EOM have a limited capacity to recover NAD^+^ independent of CI-linked mitochondrial respiration and points to a possible lack of NAD^+^ in the context of human ANT1-mediated progressive external ophthalmoplegia. Indeed, a recent study in patients with progressive external ophthalmoplegia could demonstrate an altered NAD^+^ metabolome and an improved muscle strength after boosting NAD^+^ levels using niacin (18). They further describe an increased mitochondrial biogenesis and mitochondrial respiration upon 4-months of niacin (18). In contrast, we did not observe an increased mitochondrial biogenesis, but our results suggest that the positive effect of NR is likely directly due to increased NAD^+^ availability enabling an increased glycolytic flux, as indicated by the elevated RER during running.

In summary, we demonstrate a beneficial effect of NR in mitochondrial myopathy and associated exercise intolerance in mice that is dependent on the underlying mitochondrial mutation and its effects on exercise physiology.

## Acknowledgements

We would like to thank Niagen® for providing the NR. We would like to express our gratitude to Christopher Petucci in the Metabolomics Core in the Cardiovascular Institute at the University of Pennsylvania for performing and analyzing the global and targeted metabolomics experiments. Further, we would like to thank Prof. Dr. Tejvir Khurana for his support with the sciatic nerve stimulations.

## Author contributions

P.M.S. and D.C.W. conceptualized the project and designed research; P.M.S., J.H., A.B., C.E.P., T.Y, W.T. and R.M. performed research; P.M.S. and J.H. analyzed data; P.M.S., J.A.B. and D.C.W. interpreted the data, P.M.S. and D.C.W. wrote the paper. All authors read, edited and approved the manuscript.

## Funding

This work was supported by the German Research Foundation (SCHA 2182/1-1) to PM Schaefer and National Institutes of Health grants NS021328, MH108592, OD010944 plus U.S. Department of Defense grants W81XWH-16-1-0401 and W81XWH-21-1-0128 (PR202887.e002) awarded to DC Wallace.

## Declaration of interest

Dr. Wallace is on the scientific advisory board of Pano Therapeutics and is a scientific advisor for Medical Excellence Capital. The other authors declare no conflicts of interest.

## STAR Methods Resource Availability

### Lead contact

Further information and requests for resources and reagents should be directed to and will be fulfilled by the lead contact, Dr. Douglas C. Wallace.

### Material availability

This study did not generate new unique reagents or organisms.

### Data and code availability

All data is included in the manuscript or supplementary file. Any additional information required to reanalyze the data reported in this paper in available from the lead contact upon request. Global metabolomics will be deposited at a suitable repository and made publicly available as of the date of publication.

## Experimental Model and Subject Details

### Mouse strains

All mouse strains used for this study are on the C57BL/6J with reinstated Nnt (Nnt^+/+^) background, which served as “B6 control” mice. ANT1 mice are deficient for the nuclear-encoded adenine nucleotide translocator 1 (ANT1, Slc25a4^−/−^). In addition, they harbor the complex I subunit variant ND5 m.12352C>T (ND5^S204F^). To differentiate between the effect of the ANT1 mutation and a possible effect of the ND5 variant, we created mice harboring only the mtDNA variant ND5 m.12352C>T (ND5^S204F^) (ND5 mice) as additional controls. Mice harboring the mtDNA mutation ND6 m.13997G>A (ND6^P25L^) (ND6 mice) and mice harboring the mtDNA mutation CO1m.6589T>C (CO1^V421A^) (CO1 mice) were used as additional mitochondrial mutant mouse models for the NR treatment.

Mice were fed 5LOD diet from PicoLab and kept on a 12:12 h light-dark cycle. The Institutional Animal Care and Use Committee from the Children’s Hospital of Philadelphia approved all protocols, and the protocols comply with all relevant ethical regulations regarding animal research.

## Method Details

### NR treatment

Nicotinamide riboside chloride (NR) powder was received from ChromaDex and stored light protected at 4°C. The average food consumption of mice was estimated at 3g food/24h for a 30g mouse following previous measurements in a comprehensive lab animal monitoring system. We prepared the NR diet targeting a daily dose of 400mg NR/kg body weight by mixing NR with powdered 5LOD (1:250) and water and forming pellets of 5-10g, which were air-dried for 48h under a flow hood in the dark and subsequently stored light protected at 4°C for a maximum of 7 days. Fresh NR diet pellets were provided daily (5g/mouse, 6g for ANT1 mice) and old, remaining food was removed. Food consumption was recorded for each cage containing 1-5 mice.

### Metabolic treadmill

Mice were acclimatized for 15min on the treadmill with the belt un not moving und the incline at 0°. For the exercise stress test, a stepwise ramp protocol was used. The first step at 3m/min was held for 5min Subsequently, the speed was increased every 2min to 5/7.5/10m/min continuing in steps of 2m/min. At 12/16/20/26/30 m/min the speed was maintained for 4min with the incline being increased to 5/10/15/20/25° after 2min of the respective interval. Exhaustion was defined as 5 consecutive seconds on the shock grid (0.35mA, 1shock/sec). The belt was stopped, the shock grid turned off and mice were kept in the treadmill for another 12min for recording of the recovery period. VO_2_/VCO_2_ during running at a sampling rate of 12/min with an airflow of 0.5L/min using an Oxymax (Columbus Instruments) that was calibrated once per day using a mix of 20.5% O_2_ and 0.5% CO_2_.

For hyperoxia running, the air intake was connected to an oxygen tank containing 100% oxygen. Oxygen values for were out of range for the Oxymax but VCO_2_ and running time was quantified.

### Comprehensive Lab Animal Monitoring System (CLAMS)

The Comprehensive Lab Animal Monitoring System was enclosed by an environmental chamber (Columbus Instruments) and combined with an 8-channel Oxymax system (Columbus Instruments). The system was calibrated once every 2 days using a mix of 20.5% O_2_ and 0.5% CO_2_. Air flow was set to 0.5L/min per cage.

Mice were weighted and put in the CLAMS (Columbus Instrument) during the morning (10am to noon). Mice were kept at 23°C and a 12:12 light dark cycle. Every 15min VO_2_, VCO_2_, activity, running wheel activity and food consumption were quantified starting the following day at 7am for a duration of 24h. RER (VCO_2_/VO_2_) and energy expenditure (heat=(3.815^+^1.232*RER)*VO_2_) were calculated from the data.

For the cold exposure, CLAMS were cooled down to 4°C for 4h, ending cold exposure with beginning of the light cycle at 7am.

### Rotarod

Eighteen-month-old mice were acclimatized to the rotarod prior to the first session by sitting on the non-moving rod for 1min and subsequently walking on the rod at the slowest speed (5rpm) for 1min. If mice fell, they were placed back on the rod during the acclimatization. Mice were tested for 3 consecutive days with 3 trials each day with 30min intertrial rest in their home cage. For the trials, the rod was accelerating linearly from 5rpm to 40rpm within 4min and stayed at 40rpm until the maximal test period of 10min. Time until fall was quantified, with 2 consecutive loops without intermitted walking movement equals a fall. The average across all 9 trials was calculated.

### Grip strength

18-month-old mice were grabbed by the base of their tail and allowed to hold on to a horizontal metal bar connected to a dynamometer with their front paws. They were slowly pulled away perpendicularly from the metal bar until they could not hold on anymore and the corresponding strength (N) was quantified. This was repeated consecutive 3 times in 1 session for a total of 3 sessions with 30min recovery in their home cage in between. The average grip strength across all 9 trials was calculated for each mouse.

### Lactate measurements and blood gas analysis

Immediately after exhaustive exercise (Exer 3/6 treadmill, 2m/min continuous acceleration at 10% incline for B6 control mice and 5% incline for ANT1 mice) or after sitting on the non-moving treadmill for 5min, blood was drawn using submandibular bleeding. Lactate measurements were performed using Lactate Plus Meter Test Strips (Nova biomedical). Blood gas analysis was performed using i-STAT CG8^+^ cartridges.

### Muscle pO_2_ measurements

Mice were anesthetized using 3% isoflurane in 100% oxygen and were placed on a heat pad to maintain body temperature. The gastrocnemius muscle and the sciatic nerve were uncovered and an oxygen/temperature bare fiber sensor (NX-BF/OT/E, connected to an OxyLite Pro XL) was inserted into the muscle. The sensor was minimally adjusted until a stable pO_2_ plateau was reached. Subsequently, the sciatic nerve was stimulated three times using a nerve stimulator (Grass SD9 Stimulator, 5V, 10ms pulses). The first stimulation was at 10Hz for 30sec, the 2^nd^ was at 5Hz for 1min, and the 3^rd^ was at 2.5Hz for 2min, with at least 4min recovery period between stimulations or until pO_2_ plateau was reached. The pO_2_ was continuously recorded using LabChart and the pO_2_ levels before/after stimulation as well as the average and maximal slope in the decreasing muscle pO_2_ during stimulation were quantified. Results for each stimulation frequency were combined by normalization to the baseline pO_2_ at the start of the measurement and then to the average of B6 control mice.

### Fluo-Respirometry

Mitochondrial respiration and reactive oxygen species (ROS) production in soleus muscle were assessed using the Oroboros Oxygraph-2K FluoRespirometer (Oroboros Instruments) as described previously (41). Air calibration was performed in Mir05Cr respiratory buffer [0.5 mM EGTA, 3 mM MgCl2, 60 mM lactobionic acid, 20 mM taurine, 10 mM KH2PO4, 20 mM HEPES, 110 mM D-sucrose, and fatty acid-free BSA (1 g/l) with the addition of 20 mM creatine and 5 mM DTPA] using the following settings: oxygen sensor gain: 2, gain of fluorescence-module-green: 300, data recording interval: 2 sec, block temperature: 37°C, stirrer speed: 750rpm. For ROS measurements 2µl of 10 mM AmplexTM UltraRed (Thermo Fisher Scientific), 2 µl of peroxidase (500 U/ml), 2 µl of superoxide dismutase (5 U/ml) were added to each chamber. ROS calibration was performed for each run by adding 2x 4 µl of 2.5 mM hydrogen peroxide.

Soleus muscle was dissected from mice and cut perpendicularly to allow for permeabilization, starting at the distal end. Two muscle pieces were weighed, suspended in Mir05Cr respiratory buffer and added into a respiration chamber in 400 µl of Mir05Cr (performed in technical duplicates).

Subsequent addition of substrates allowed assessment of different respiratory states. First, 5 µl of 2 M pyruvate, 2.5 µl of 400 mM malate, and 10µl of 2 M glutamate (PMG) were added, followed by 20 µl of 0.5 M adenosine diphosphate to provide complex I respiration. Addition of 20µl of 1 M succinate results in OxPhos capacity. Increase of respiration with succinate also validated that complex I respiration is not limited by the mitochondrial coupling. 0.5 µl of 10 mM oligomycin blocked complex V and provides Leak respiration, followed by titration of the uncoupler carbonyl cyanide-p-trifluoromethoxyphenylhydrazone (FCCP, 1 mM stock in 1 µl increments) to reveal the electron transport system capacity (ETS capacity). 5 µl of 1 mM rotenone was added, resulting in complex II respiration. Finally, 2 µl of 5 mM antimycin A blocked CIII, providing non-mitochondrial background respiration, which was subtracted from all other respiratory states. Reoxygenation of the chambers at the end of the titration protocol allowed a quantification of ROS production at the same respiratory state with 2 different oxygen concentrations. The resulting equation was used to correct ROS production for the effect of O2 concentration in the chamber. In addition, ROS production in the presence of all uncouplers and inhibitors provides a surrogate marker for mitochondrial mass. Data were recorded and analyzed using DatLab7. Complex I and II capacities were calculated as maximum of (coupled respiration with PMG or ETS capacity minus complex II respiration) for complex I and (OxPhos capacity minus coupled PMG or uncoupled respiration after rotenone) for complex II.

### Fluorescence lifetime imaging microscopy (FLIM) of NADH

Immediately after cervical dislocation of a mouse, one gastrocnemius muscle was dissected and placed in Tyrodes buffer. A muscle bundle was separated carefully in Tyrodes buffer and placed on an LSM710 (Zeiss) in an imaging chamber (Warner Instruments) in Tyrodes buffer. Imaging was performed at room temperature within 30min of preparation using a 20x lens and pulsed, 2-photon excitation (730nm, 80MHz, 10fs pulse width, 5mW power on sample, 1min excitation time) of NADH. NADH autofluorescence lifetime was detected through a 460/50nm bandpass filter using time-correlated single photon counting (HPM-100-40, Becker&Hickl) with a time resolution of 256 time channels within the 12.5ns pulse period (512×512 pixels, 15µsec pixel dwell time, SPCM 9.8). A biexponential decay with lifetime components fixed to 400ps (free NADH) and 2500ps (protein-bound NADH) was fitted to the NADH autofluorescence decay curve for every pixel (bin 2) using SPCImage 8.0 and the mean NADH lifetime (τ_mean_) was quantified for every image.

### Global metabolomics

After exhaustive exercise (Exer 3/6 treadmill, 1m/min continuous acceleration at 10% incline for B6 control mice and 5% incline for ANT1 mice) or after sitting on the non-moving treadmill for 5min mice were cervically dislocated and the gastrocnemius muscle was dissected first and immediately flash-frozen in liquid nitrogen and stored at −80°C.

### Untargeted LC/MS Metabolomics

Wet, frozen mouse gastrocnemius samples were lyophilized overnight and powdered in a Precellys homogenizer (Bertin Technologies) at 4 °C. Approximately 5 mg aliquots of dry tissue powder were homogenized in 500 µL of 80% ice-cold methanol in the Precellys homogenizer at 4 °C. Then, 100 µL of homogenates were extracted with 400 µL of ice-cold methanol, vortexed, centrifuged at 18,000 x g, and 400 µL aliquots of supernatants were dried under nitrogen at 45 °C in a 96-well plate. Samples were reconstituted in 10% methanol for reversed-phase C18 chromatography or 50% methanol for HILIC chromatography with gradient elution. In addition, 10 µL of each homogenate was pooled together, divided into 100 µL aliquots, and extracted according to the above protocol to make QC samples. Samples were analyzed by a Thermo Vanquish UHPLC/Orbitrap ID-X mass spectrometer scanned from *m/z* 60-1000 at a resolution of 120,000. Compound Discoverer (Thermo Fisher Scientific) was used to generate PCA plots from the metabolite signals extracted from the raw data files, fold changes, p-values, heat maps, whisker plots, and perform a database search for metabolite identification. Pathway analysis was performed using Metascape (44).

### Targeted LC/MS Metabolomics

Approximately 5 mg aliquots of dry gastrocnemius powder were homogenized in 500 µL of 50% acetonitrile/0.3% formic acid for the extraction of organic acids and malonyl and acetyl CoA. Additional 5 mg aliquots of dry tissue powder were homogenized in 500 µL of 80% methanol for the extraction of nucleotides in a similar way to the untargeted metabolomics protocol above. Aliquots (100 µL) of the homogenates prepared for organic acids and malonyl and acetyl CoA were extracted and quantitated by LC/MS according to validated, optimized protocols in our previously published studies (42, 43). Nucleotides were extracted from 100 µL homogenates with 400 µL of ice-cold methanol and processed further according to the untargeted metabolomics protocol above prior to LC/MS. An Agilent PEEK poroshell HIILIC-z column was used to separate nucleotides with gradient elution. Quantitation of metabolites in each assay module was achieved using multiple reaction monitoring of calibration solutions and study samples on an Agilent 1290 Infinity UHPLC/6495 triple quadrupole mass spectrometer. Raw data were processed using Mass Hunter quantitative analysis software (Agilent). Calibration curves (R2 = 0.99 or greater) were either fitted with a linear or a quadratic curve with a 1/X or 1/X^2^ weighting.

### NAD/ NADH quantification

For quantification of NAD and NADH in gastrocnemius muscle, we powdered flash-frozen gastrocnemius on dry ice, split the powder into 4 tubes, quantified the weight, flash froze the gastrocnemius powder in liquid nitrogen and stored at −80°C.

Two tubes were used for the NAD/NADH cycling assay. For NAD, powdered tissue was lysed in ice-cold 0.6M Perchloric Acid, for NADH in ice-cold 0.25M KOH in 50% ethanol and spun down for 15min at 20,000g at 4°C. Supernatant is transferred to a new tube and diluted 1:40 (1:25 for NADH) in ice cold Na-Phosphate buffer (pH 8.0). NAD standards or diluted extracts were added to a cycling mixture consisting of 2% ethanol, 100 μg/mL alcohol dehydrogenase, 10 μg/mL diaphorase, 20 μM resazurin, 10 μM flavin mononucleotide, 10 mM nicotinamide, 0.1% BSA in 100 mM phosphate buffer, pH 8.0. Fluorescence was quantified at 590nm (ex:530nm) in a 96 well plate.

One tube was used for NAD/NADH quantification using the NAD/NADH Glo™ Assay (Promega) following the manufacturer’s protocol. The results of both assays were normalized to the average across all samples and subsequently combined for each mouse.

## Quantification and Statistical Analysis

In all scatter plots, each data point represents one independent mouse with the mean and standard error of the mean (SEM) being indicated by the error bars. The number of independent experiments and the respective statistical tests are indicated in the figure legends. Statistical analysis was performed using Graph Pad Prism 5. For VO_2_ and RER kinetics during the exercise stress test, the average of all active mice is displayed until half of the mice stopped. For in-vivo pO_2_ measurements (Fig.2e, f) box plots with whiskers 10-90% are used for improved clarity due to a high variance. Independent experiments for respirometry are defined as independent individual mice sacrificed and measured in technical duplicates, and their average (mean) is used as the independent data point. For microscopy, independent measurements (data points) were defined as images from individual mice with 5 images averaged per mouse. For metabolomics, independent experiments indicate tissue samples from individual mice. Statistical analysis for metabolomics are described in the methods section.

## Supplemental Information

### Supplementary Figure Legends

**Figure S1:**
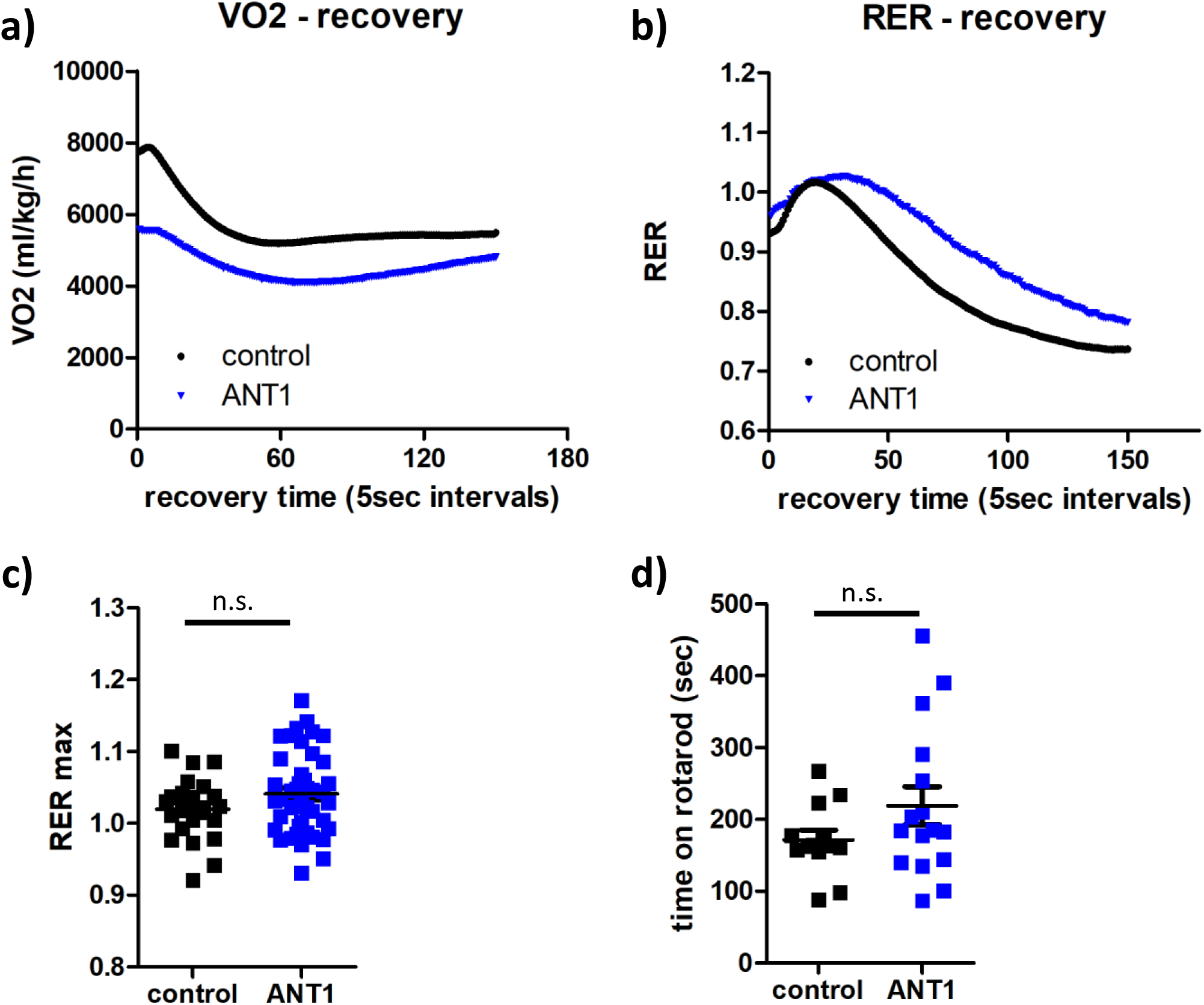
ANT1 mice show no neurological limitation of exercise capacity. **a-c)** Kinetics of the average oxygen uptake (VO_2_) (a), respiratory exchange ratio (RER, VO_2_/VCO_2_) (b) and maximal RER (c) of 4-month-old mice after an exercise stress test (ramp protocol) on a metabolic treadmill (n = 24-40). d) Average time of 18-month-old B6 control and ANT1 mice on an accelerating rotarod until non-compliance (n = 13-16, average of 9 trials/mouse). Significances between strains (*) were calculated using unpaired t-test (*p < 0.05, **p < 0.01, ***p < 0.0001).

**Figure S2:**
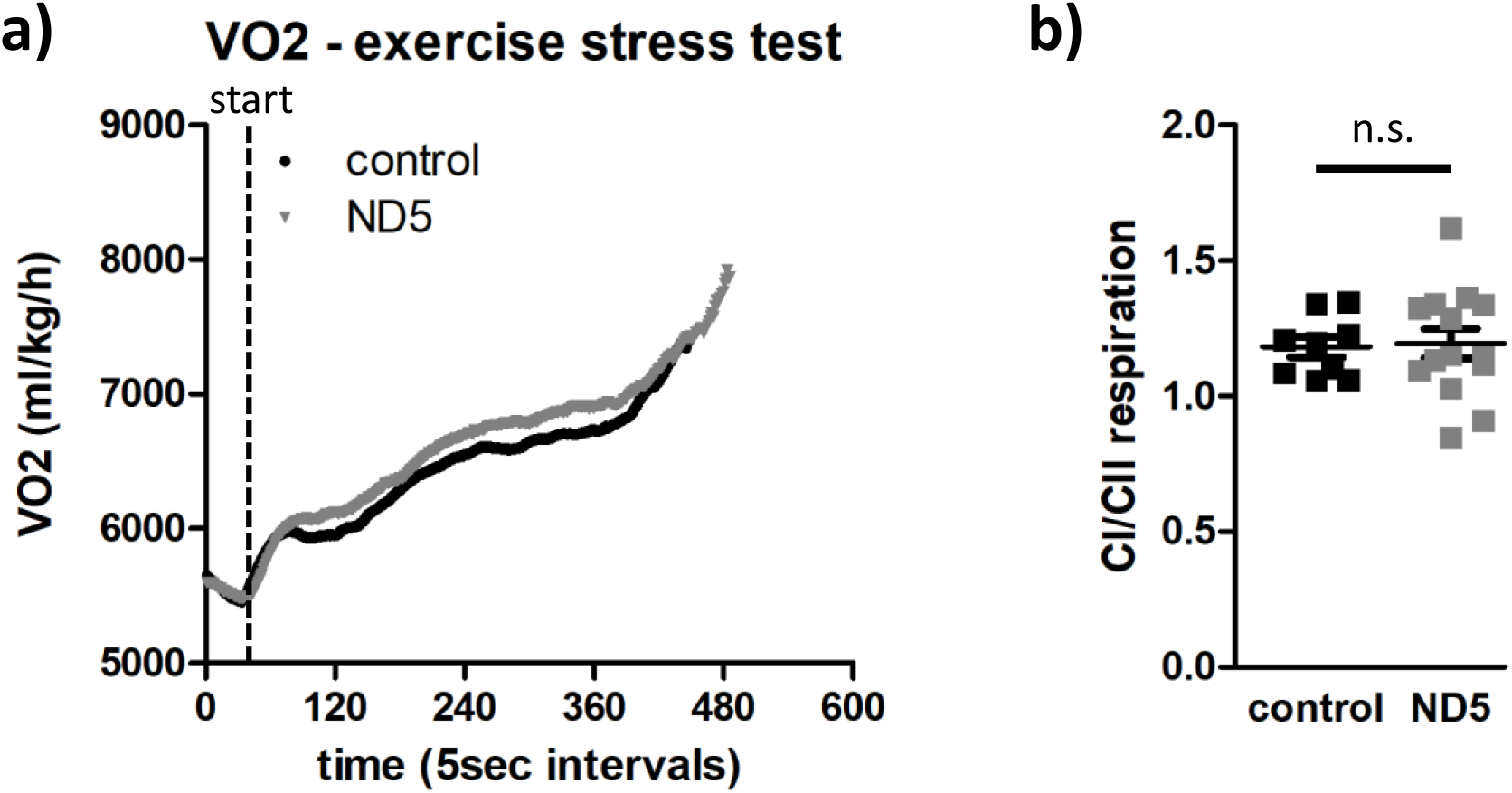
ND5 mutation does not affect exercise capacity or CI respiration. **a)** Kinetics of the average oxygen uptake (VO_2_) of 4-month-old B6 control or ND5 mice after an exercise stress test (ramp protocol) on a metabolic treadmill (n = 24-40, curve displayed until >50% of the animals quit). **b)** CI/CII respirational capacity of M. soleus in 6-month-old B6 control and ND5 mice. Significance between strains (*) was calculated using unpaired t-test. (*p < 0.05).

**Figure S3:**
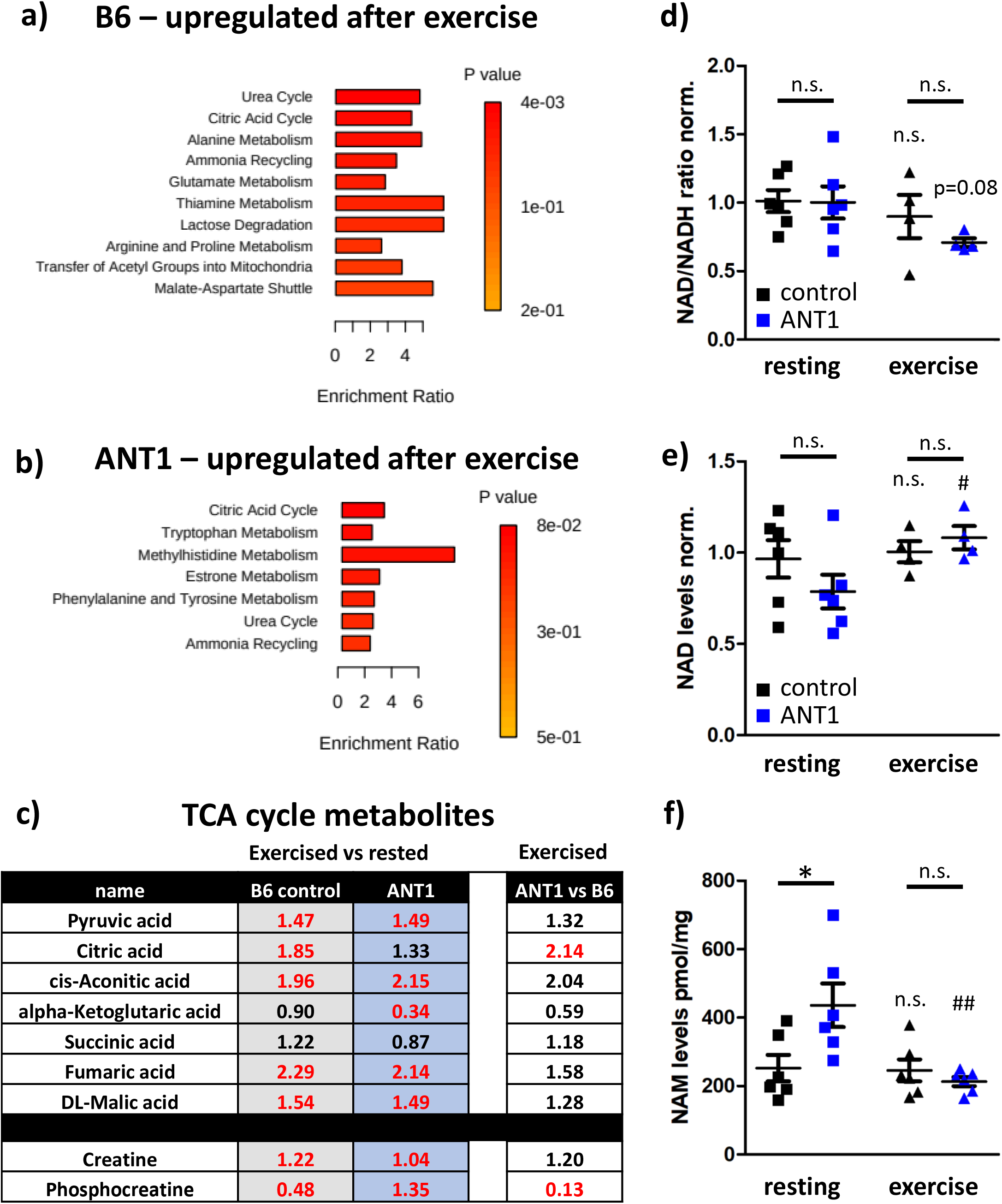
Metabolomics in skeletal muscle after exercise. **a/b)** Enrichment analysis of global metabolomics (upregulated metabolites, p < 0.05) in M. gastrocnemius of resting B6 control (a) and ANT1 (b) mice compared to mice exercised until exhaustion using Metascape. The enrichment ratio is indicated by the length of the bar and the p-value by color. **c)** Fold changes of TCA cycle metabolites and creatine levels in exercised vs rested B6 controls (column 2), exercised vs rested ANT1 mice (column 3) and between exercised ANT1 vs exercised B6 control mice (column 4). Fold changes in red font indicate a significant change in metabolite levels, black is non-significant. **d/e)** NAD^+^/NADH ratio (d) and NAD levels (e) in M. gastrocnemius normalized to resting B6 control mice (n = 4-6). **f)** Nicotinamide (NAM) levels in M. gastrocnemius of resting mice and immediately after exhaustive exercise (n = 6). Significances between strains (*) and between resting and exercise (#) were calculated using Mann Whitney or unpaired t-test (*p < 0.05, **p < 0.01, ***p < 0.0001).

**Figure S4:**
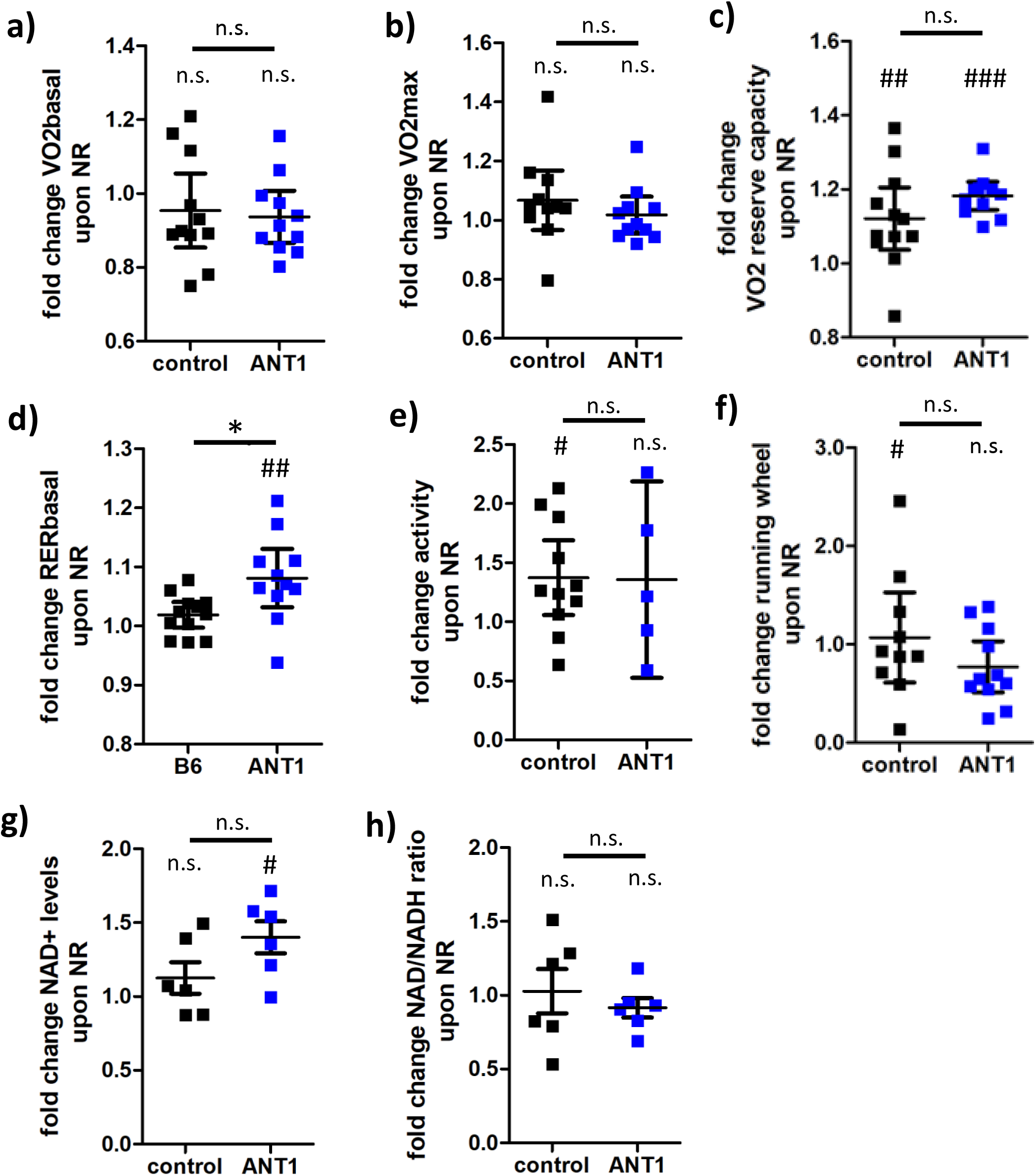
Nicotinamide riboside improves exercise physiology in ANT1 mice. **a/b/c/d)** Fold change in VO_2_ during rest (a), VO_2_max (b), VO_2_ reserve capacity (VO_2_basal/VO_2_max) (c), and RER during rest (d) in an exercise stress test upon an 8-week treatment with NR in 6-month-old B6 control and ANT1 mice (n = 11-12). **e/f)** Fold change in activity levels (e) and running wheel activity (f) during 24h spent in CLAMS upon an 8-week treatment with NR in 6-month-old B6 control and ANT1 mice (n = 5-12). **g/h)** Fold change in NAD levels (g) and NAD/NADH ratio (h) upon an 8-week treatment with NR in 6-month-old B6 control and ANT1 mice (n = 6). Significances between strains (*) and between NR-treated and untreated (#) were calculated using Mann Whitney or unpaired t-test (*p < 0.05, **p < 0.01, ***p < 0.0001).

